# Rapid cortical plasticity induced by active associative learning of novel words in human adults

**DOI:** 10.1101/716712

**Authors:** Alexandra M. Razorenova, Boris V. Chernyshev, Anastasia Yu. Nikolaeva, Anna V. Butorina, Andrey O. Prokofyev, Nikita B. Tyulenev, Tatiana A. Stroganova

## Abstract

Human speech requires that new words are routinely memorized, yet neurocognitive mechanisms of such acquisition of memory remain highly debatable. Major controversy concerns the question whether cortical plasticity related to word learning occurs in neocortical speech-related areas immediately after learning, or neocortical plasticity emerges only on the second day after a prolonged time required for consolidation after learning. The functional spatiotemporal pattern of cortical activity related to such learning also remains largely unknown. In order to address these questions, we examined magnetoencephalographic responses elicited in the cerebral cortex by passive presentations of eight novel pseudowords before and immediately after an operant conditioning task. This associative procedure forced participants to perform an active search for unique meaning of four pseudowords that referred to movements of left and right hands and feet. The other four pseudowords did not require any movement and thus were not associated with any meaning. Familiarization with novel pseudowords led to a bilateral repetition suppression of cortical responses to them; the effect started before or around the uniqueness point and lasted for more than 500 ms. After learning, response amplitude to pseudowords that acquired meaning was greater compared with response amplitude to pseudowords that were not assigned meaning; the effect was significant within 144–362 ms after the uniqueness point, and it was found only in the left hemisphere. Within this time interval, a learning-related selective response initially emerged in cortical areas surrounding the Sylvian fissure: anterior superior temporal sulcus, ventral premotor cortex, the anterior part of intraparietal sulcus and insula. Later within this interval, activation additionally spread to more anterior higher-tier brain regions, and reached the left temporal pole and the triangular part of the left inferior frontal gyrus extending to its orbital part. Altogether, current findings evidence rapid plastic changes in cortical representations of meaningful auditory word-forms occurring almost immediately after learning. Additionally, our results suggest that familiarization resulting from stimulus repetition and semantic acquisition resulting from an active learning procedure have separable effects on cortical activity.

## 1 Introduction

Words are distinct, meaningful elements of a human language. Recognition of spoken words requires the brain to possess phonological representations of complex auditory patterns that represent words (Griffiths and Warren, 2004; DeWitt and Rauschecker, 2012). Words also bear meaning which allows using them as information carriers for inter-subject communication (Fodor, 1983). However, it remains poorly understood how learning new words is implemented in the brain. Major controversy concerns the question whether cortical effects of lexical acquisition need a prolonged time for consolidation. Also, precise timing, latency, and localization of learning effects in the brain are poorly understood.

The two-stage complementary learning systems theory (Davis and Gaskell, 2009), based mainly on functional magnetic resonance imaging (fMRI) and positron emission tomography (PET) evidence, posits that formation of a new word representation, similarly to formation of other long-term memory traces, is a two-stage process (Gaskell and Dumay, 2003; Davis and Gaskell, 2009; Rodríguez-Fornells et al., 2009). According to this theory, it initially involves rapid but short-lived learning of a new word, mainly subserved by the medial temporal memory system without substantial neocortical involvement. The slowly emerging plastic changes in neocortical responses occur through off-line consolidation, i.e., strengthening of word representation within neocortical networks, and presumably require an overnight sleep. Accordingly, it could be unlikely to find manifestations of cortical plasticity on the first day of lexical learning.

Yet, in recent years, research has been made that suggests a faster mechanism of neocortical word plasticity that does not involve a prolonged period of consolidation (Mestres-Missé et al., 2007; Shtyrov et al., 2010; Sharon et al., 2011; Borovsky et al., 2013; Kimppa et al., 2015; Hebscher et al., 2019).

Rapid neocortical plasticity during word acquisition has been addressed in a large body of electroencephalographic/magnetoencephalographic (EEG/MEG) studies, which sought evidence of fast and automatic formation of phonological word-form cortical representations that results from mere repetitive presentations of pseudowords during passive listening. Such data demonstrated that while a cortical response to pseudowords was initially weaker than that to real words, after a number of repetitions the response to novel pseudowords was increased and the difference was diminished (Shtyrov et al., 2010; Shtyrov, 2011; Kimppa et al., 2015; Kimppa et al., 2016; Partanen et al., 2017). Similar results were also obtained in mismatch-negativity (MMN) studies (Yue et al., 2014; Yue et al., 2017; Kompus and Westerhausen, 2018). These findings were interpreted as evidencing that the adult cerebral neocortex brain can learn novel pseudowords in the course of passive listening without any cognitive or attentional effort, and, importantly, the effect could be obtained within several tens of minutes. Yet, these EEG and MEG studies addressed only the phonological aspect of lexicality, while a semantic aspect was beyond their scope. Potentially, such rapid phonological plasticity could be explained by a more general biological mechanism of implicit perceptual learning (Seitz and Dinse, 2007) rather than by specific mechanisms of linguistic learning. For example, a mere familiarization with unattended nonverbal visual stimuli could lead to improved discrimination (Sasaki et al., 2010). While EEG/MEG studies mentioned above used phonological learning thus addressing only one aspect of lexicality acquisition, the most common approach in fMRI research (Rodríguez-Fornells et al., 2009) relied upon associative procedures to address both phonological and semantical learning.

To the best of our knowledge, there were only few EEG studies dedicated to associative learning of novel auditory words, addressing phonological and semantic learning jointly. Associative learning procedures used in the field were intended to establish a link between novel word-forms (pseudowords) and referents carrying meaning. Most usually, associative learning procedures involved paired presentations of auditory novel pseudowords and their intended referents such as visual images, movies or real words presented in conjunction with pseudowords. Anyway, the associative procedure in most of such studies was passive in the sense that no active response choice was expected from participants during the learning phase. In one of such EEG studies (Fargier et al., 2014), pseudowords were associated either with short movies of reaching-and-grasping movements or with abstract visual images. Event-related potentials (ERPs) were compared to passive pseudoword presentation before and after learning. After learning, ERP started to differentiate both types of pseudowords within 100 – 400 ms after stimulus onset. In line with fMRI findings and the two-stage complementary learning systems theory (Davis and Gaskell, 2009), reliable learning-induced changes in ERPs occurred only on the second day after learning, supposedly after an overnight consolidation. Thus, this ERP study provided no confirmation for rapid semantic cortical plasticity in word learning.

The other available EEG study (François et al., 2017) explored efficacy of associative learning in comparison with statistical learning when the participants learned four tri-syllabic pseudowords presented within a continuous stream of auditory syllables. The results showed that during the learning phase, the “semantic” N400 component of the ERP (Kutas and Federmeier, 2011) was elicited by pseudowords associated with visual images but not by control pseudowords, thus bringing evidence in favor of fast semantic cortical plasticity. However, the findings obtained in these ERP studies may be difficult to interpret unambiguously because the effect was measured during the learning blocks of the experiment. Indeed, since participants were required to listen carefully to the auditory stream with the task of discovering new words, learning-related enhancement of N400 might have been elicited by attention biased towards auditory word-forms that were accompanied by pictures. Ideally, in order to prove semantic cortical plasticity, the effect should be probed during passive exposures to newly learned pseudowords after learning in comparison with identical exposures before learning.

Quite recently, a study was published that overcame a limitation mentioned above by measuring MMN during passive sessions before and after associative learning rather than during learning (Aleksandrov et al., 2019). As a result of training, the amplitude of the MMN to novel word-forms was enhanced. The associative learning procedure was also passive and involved simple pairing of a novel word-form with a real word used as a referent in this study. Limitations of this study included a very small number of novel pseudowords and corresponding referent items (just two), and a non-counterbalanced procedure.

In contrast to the EEG studies mentioned above that used a passive learning procedure, an active learning procedure was used by Hawkins et al. (2015), who found increased MMN in response to auditory pseudowords that acquired meaning through association with visual images, while no effect was detected for similar word-forms that were not consistently associated with any referents. The effect was present during a passive oddball session administered immediately after an active learning procedure. Surprisingly, on the second day, after a 24-hour consolidation period, the enhanced MMN effect disappeared, while auditory discrimination was still evident behaviorally. None of the mentioned EEG studies that used associative procedures attempted to localize brain sources.

Another EEG study demonstrated ERP enhancement recorded during passive presentations before and after a fast mapping procedure (Vasilyeva et al., 2019). Maximal effect was observed in the left temporal cortex and in the left anterior prefrontal cortex, yet the authors acknowledged that source localization had technical limitations and should be treated with caution. Notably, fast mapping is a process of exclusion-based inference (Carey and Bartlett, 1978); thud it involves an active decision and response selection on the part of the participant.

Altogether, at least some of the EEG studies mentioned above contradict the two-stage complementary learning systems theory (Davis and Gaskell, 2009) by evidencing cortical effects of novel word-form learning without an overnight consolidation; due to the nature of the EEG signal, effects in medial temporal locations including the hippocampus could not be detected in these studies.

It should be noted that at a few MRI studies also reported neocortical effects immediately after learning, in addition to changes in activity within the hippocampus, which is typically observed in MRI studies of word acquisition. Importantly, both studies used active association procedure similar to operant conditioning. The effects found in one of them (Breitenstein et al., 2005) involved the left inferior parietal cortex, while no effects were reported for other neocortical speech areas. The other study reported significant microstructural changes in a much wider set of cortical regions involved in language and reading, including inferior frontal gyrus, middle temporal gyrus, and inferior parietal lobule (Hofstetter et al., 2017). Imaging sessions in this study were done before and immediately after the learning task.

In summary, the current picture of fast semantic cortical plasticity in word learning is far from complete. Fast effects related to word learning were detected in a large number of EEG studies (Shtyrov et al., 2010; Shtyrov, 2011; Yue et al., 2014; Kimppa et al., 2015; Kimppa et al., 2016; Partanen et al., 2017; Yue et al., 2017; Kompus and Westerhausen, 2018). Yet the passive nature of learning procedures used in most previous experiments causes some skepticism. A word, which is learned passively through repetition or instructions, is typically not well retained or effectively used. Active search for word meaning might be a preferred mode for inducing fast semantic learning. Indeed, animal data suggest that the most effective way to induce cortical plasticity in adult primates is the operant conditioning paradigm. For example, a series of studies by Blake and colleagues (Blake et al., 2002; Blake et al., 2005; Blake et al., 2006) showed that a fast and permanent transformation in cortical neuronal activity occurs in primates only if an active operant conditioning procedure is used (and not through passive stimulus-reward associative pairing). The neuroimaging studies that used active learning procedures similar to operant conditioning (Breitenstein et al., 2005; Hawkins et al., 2015; Hofstetter et al., 2017) or fast mapping procedures (Merhav et al., 2015; Vasilyeva et al., 2019) are very promising in this respect.

A theoretical framework has recently been put forward to support the mechanisms of rapid cortical plasticity (Hebscher et al., 2019); specifically an active learning procedure has been ascertained as a prerequisite condition for such cortical plasticity. Additionally, there is still no reliable data on localization of cortical plasticity in time resolved EEG/MEG studies, while fMRI studies cannot provide exact timing of the effects. And there remains a profound ambiguity on the relation between familiarization and sematic learning as likely constituent parts of lexicalization.

In the current MEG study, which employed an active operant conditioning task, we sought evidence of putative rapid cortical plasticity linked to two interrelated yet separate processes: formation of a new acoustic word-form discrimination and semantic analysis of the newly-formed word item. To pursue this goal, we engaged our participants in an associative learning task to let them actively find unique associations between four auditory pseudowords and their own body part movements, whereas the other four auditory pseudowords were not associated with any motor action. To reveal the learning effect, we compared responses to passive presentations of the same auditory pseudoword stimuli immediately before and immediately after the learning sessions. We used MEG neuroimaging technique, which offers the best combination of excellent time resolution and good spatial resolution. These factors allowed us to identify the anticipated effects both in terms of their timing and cortical regions involved. In contrast to the previous EEG/MEG studies aiming at rapid cortical plasticity, we did not focus our analysis on pre-specified cortical regions or the time intervals of cortical responses, and we employed an unbiased data-driven search (with correction for multiple comparisons) to reveal when and where in the cortex learning of novel word-forms and/or acquiring their semantics would induce neural activity changes.

We hypothesized that we would find evidence of rapid cortical plasticity immediately after learning. We expected this especially with the use of an active associative task, which may promote strong transformation of cortical activity following associative learning (Blake et al., 2002; Blake et al., 2005; Blake et al., 2006). We expected that the effects of familiarization and semantization would be separable. We hypothesized that changes in brain activity caused by word-form familiarization would be observed both for pseudowords that acquire a unique association with a specific movement and for those that do not. We expected to see these respective changes in time-locked cortical responses rather early (starting ~100–200 ms after pseudo-word onset) in the perisylvian speech areas, which are thought to be engaged in phonological processing of an auditory word-forms (DeWitt and Rauschecker, 2012). Secondly, we expected to find the cortical plasticity signature in the semantic brain network. We anticipated finding modulation of cortical activity by learning at a later time in the higher-tier speech areas in the temporal and frontal cortices that mediate semantic analysis of wordforms (Davis and Gaskell, 2009; Kutas and Federmeier, 2011). Critical for our hypothesis, we predicted that the latter “semantic” modulation would be observed selectively for the meaning-related pseudowords and would be absent for the well-familiarized but meaningless pseudowords.

## 2 Materials and Methods

### 2.1 Participants

Twenty-four volunteers (mean age 24.9 years, range 19–33 years, 15 males) participated in the study. They were native Russian speakers with normal hearing and no record of neurological or psychiatric disorders. All participants were right-handed according to the Edinburgh Handedness Inventory (Oldfield, 1971). The study was conducted following the ethical principles regarding human experimentation (Helsinki Declaration) and approved by the Ethics Committee of the Moscow State University of Psychology and Education. All participants signed the informed consent before the experiment.

### 2.2 Stimuli and behavioral responses

The auditory stimuli (pseudowords) were created in such a way to precisely control and balance their acoustic and phonetic properties while manipulating their lexical status before and after learning. We used nine consonant-vowel (CV) syllables, which formed eight disyllabic (C_1_V_1_C_2_V_2_) novel meaningless word-forms (pseudowords). The pseudowords were built in compliance with Russian language phonetics and phonotactic constraints. After the associative learning procedure, four of them were assigned a unique action performed by one of four body extremities (action pseudowords – APW), while the other four implied no motor response (non-action pseudowords – NPW).

The first two phonemes (C_1_V_1_) formed the syllable ‘hi’ [x^J’^i] that was identical for all pseudowords used. The next two phonemes (C_2_ and V_2_) were independently counterbalanced across APW and NPW stimuli, and they were included in the two stimuli of each type, forming eight unique phonemic combinations (Table 1). This design ensured that acoustic and phonetic features were fully matched between the APW and NPW types (within respective pairs). The third phonemes (C_2_), consonants *‘ch’* 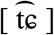, *‘sh’* [ʂ], *‘s’* 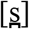, *‘v’* [v], distinguished between the APW–NPW pairs by signaling which extremity a subject might be prepared to use (right hand, left hand, right foot, or left foot). All of the pseudo-words could only be recognized by their fourth phoneme (V_2_: vowel *‘a’* [ə] or *‘u’* [ʊ]). The onset of the fourth phoneme will be referred to as “word-form uniqueness point” (UP; Figure 1A).

**Table 1.**
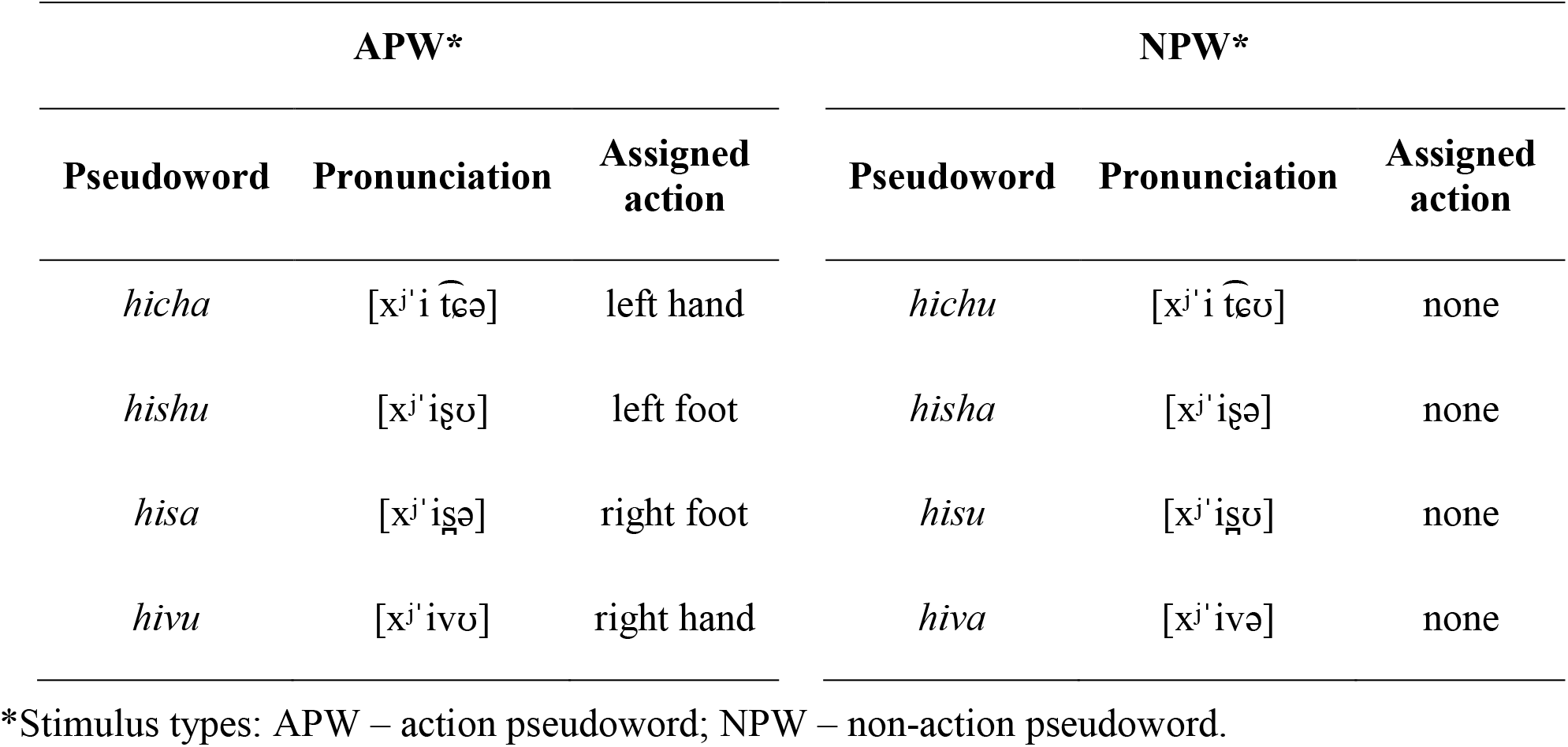
Stimulus-to-response mapping.

**Figure 1.**
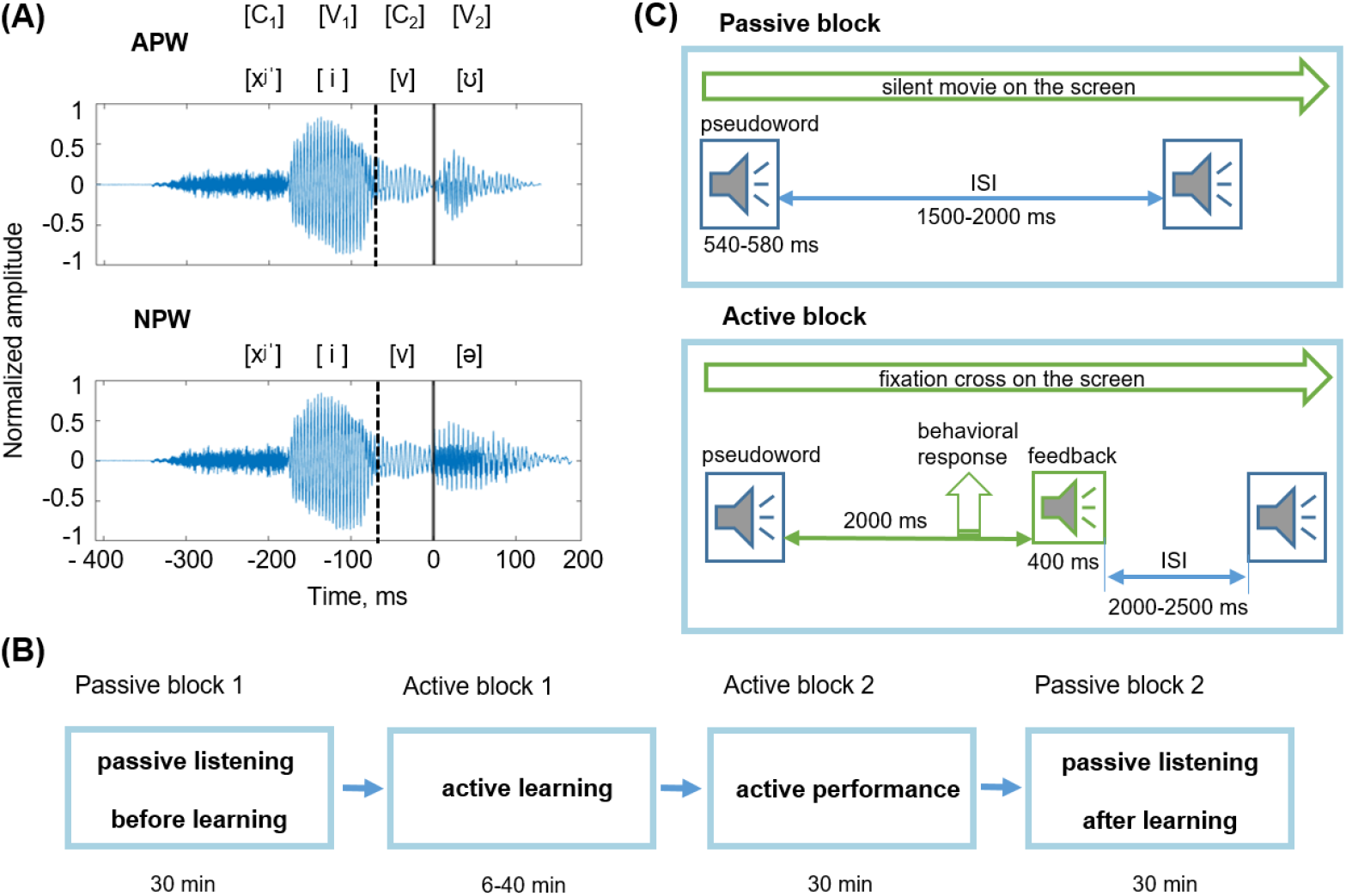
Stimuli and experimental design. **(A)** Examples of pseudoword stimuli: ‘hivu’ and ‘hiva’. All stimuli were two-syllable pseudowords (C_1_V_1_C_2_V_2_). The first syllable C_1_V_1_ (‘hi’) was the same for all pseudowords. Pseudowords were organized in pairs; each pair differed from the other pairs by the third phoneme, the consonant C_2_. Each pair included an action pseudoword (APW) and a non-action pseudoword (NPW), which differed from each other by the last vowel V_2_ (either ‘a’ or ‘u’; Table 1). Here and hereafter, a zero value on a timeline and a vertical solid line denote the onset of the fourth phoneme (word-form uniqueness point [UP]); a vertical dashed line indicates the onset of the third phoneme. **(B)** The sequence of experimental blocks. **(C)** The experimental procedure during passive blocks (upper panel) and active blocks (bottom panel); ISI refers to the interstimulus interval. During both passive blocks, participants were offered to watch a silent movie while auditory stimuli were presented. During active blocks, participants learned associations between pseudowords and motor actions.

As can be seen in Table 1, the phonetic composition of the stimuli and stimulus-to-response mapping complied with a full within-subject counterbalanced design, in relation to the third and fourth phonemes, as well as in relation to movements by left/right and upper/lower extremities.

All stimuli were digital recordings (PCM, 32 bit, 22050 Hz, 1 channel, 352 kbps) of a female native Russian speaker’s voice recorded in a sound-attenuated booth. Four variants of three-phoneme combinations (C_1_V_1_C_2_) and two variants of the last vowel (V_2_) were recorded and then combined to generate eight pseudowords. All pseudowords were pronounced with stress on the vowel ‘i’ in order to match prosody between all the utilized pseudowords. The amplitude of the recorded stimuli was digitally equalized by maximal power, which corresponded to the stressed vowel ‘i’. For cross-splicing and normalization, sound recordings of the pseudowords were digitally processed using Adobe Audition CS6.5 software. Duration of the pseudowords was approximately 530 ms. For all analyses, data were aligned on the word UP, which was kept at 410 ms after the onset of the audio recordings.

Additionally, two non-speech auditory stimuli were used as positive and negative feedback signals, each 400 ms in length. Both stimuli were complex frequency-modulated sounds that profoundly differed in their spectral frequency maxima (ranges were approximately 400–800 Hz for positive and 65–100 Hz for negative feedback), with spectral maxima increasing in frequency over time for the positive feedback and decreasing for the negative feedback.

Behavioral responses (Table 1) were recorded using hand-held buttons (package 932, CurrentDesigns, Philadelphia, PA, USA) pressed by the right or left thumb and custom-made pedals pushed by the toes of the right or left foot. For all of these movements, the actual trajectory was rather short (< 1 cm for buttons and < 3 cm for pedals), a design that minimized movement artifacts. Buttons and pedals interrupted a laser light beam delivered via fiber optic cable. Responses recorded from pedals and buttons were automatically labeled as ‘correct’ and ‘errors’ after each trial according to the task rules (see below).

### 2.3 Procedure

During the experiments, participants were comfortably seated in the MEG apparatus that was placed in an electromagnetically and acoustically shielded room (see below). Pseudowords were presented binaurally via plastic ear tubes in an interleaved quasi-random order, at 60 dB SPL. The experiment was implemented using the Presentation 14.4 software (Neurobehavioral systems, Inc., Albany, CA, USA).

The experiment consisted of four consecutive blocks with a fixed order across participants: (1) passive listening before learning, (2) active learning, (3) active performance, and (4) passive listening after learning (Figure 1B). The entire experiment lasted approximately 2 hours.

Two identical passive listening blocks were administered before and after the two active blocks. During passive auditory presentation, participants were offered to watch a silent movie projected on the screen positioned at eye-level 2 m away. Pseudowords were presented pseudo-randomly with an average interstimulus interval (ISI) of 1750 ms, randomly jittered between 1500 and 2000 ms at 1 ms steps (Figure 1C). Each passive listening block included 400 stimuli (50 repeated presentations of each of eight pseudowords) and lasted approximately 30 min.

After the first passive block, the participants were informed that during the following active blocks they had to find the association between each of the presented eight pseudowords and movements of their own body parts. To achieve this goal, they were asked to respond to each pseudoword either by using one of the four body extremities or to commit no response, and then to listen to positive and negative feedback signals informing the participants whether the action was correct or erroneous. The instruction did not contain any other cues. The behavioral procedure used, which involved trying a variety of new auditory-action associations and eventually selecting only those that led to positive reinforcement, complied with the requirements of operant learning (Neuringer, 2002).

During the active learning block, participants were required to keep their gaze at the fixation cross in the center of the presentation screen in order to minimize artifacts caused by the participants’ eye movements. The eight pseudowords were repeatedly presented within pseudo-random interleaved sequences. For each trial, a pseudoword was followed by a feedback signal, which was presented 2000 ms after the end of the pseudoword stimulus (Figure 1C). The average ISI (from the end of the feedback stimulus until the onset of the next pseudoword stimulus) was 2250 ms, randomly jittered between 2000 and 2500 ms at 1 ms steps. The feedback stimulus could be either positive or negative. Positive feedback was given if a participant complied with the task rues, i.e., committed a proper response to an APW stimulus or committed no response to an NPW stimulus (Table 1). Negative feedback followed three kinds of errors: (i) no response to an APW; (ii) a motor response to an APW performed with “a wrong extremity”; (iii) any response to an NPW. The number of stimuli in this block varied across participants depending on the individual success rate. An active learning block ended if a participant reached the learning criterion or if 480 stimuli were presented in total, whichever came first. Successful learning implied that a participant performed the correct responses in at least four out of five consecutive repeated presentations of each of the eight pseudowords.

Whether a participant met the learning criterion was automatically checked after each trial. Out of 24 participants, two did not reach the learning criterion and thus went through all 480 trials in the learning block. Since their overall hit rate during the next active performance block was well within the range of performance of the other 22 participants, these two participants were not excluded from further analyses. The number of stimuli presented within the active learning block varied across participants from 74 to 480, with the respective inter-individual variation in the duration of active learning from 6 to 40 min.

Participants were then asked to repeat the same procedure (active performance block). The only difference between the two active blocks was that the active performance block included a fixed number of 320 trials and lasted approximately 30 min.

Short breaks were introduced between all blocks (10 min between the active performance block and the second passive block and 3 min between other blocks), during which participants were offered to rest while remaining seated in the MEG apparatus.

### 2.4 MEG data acquisition

MEG data were recorded inside a magnetically shielded room (AK3b, Vacuumschmelze GmbH, Hanau, Germany), using a dc-SQUID Neuromag VectorView system (Elekta-Neuromag, Helsinki, Finland) with 204 planar gradiometers and 102 magnetometers. For all recorded signals, the sampling rate was 1000 Hz, and the passband was 0.03–330 Hz.

Participants’ head shapes were measured using a 3Space Isotrack II System (Fastrak Polhemus, Colchester, VA, USA) by digitizing three anatomical landmark points (nasion, and left and right preauricular points) and additional randomly distributed points on the scalp. During MEG recording, the position and orientation of the head were continuously monitored by four Head Position Indicator coils.

The electrooculogram was registered with two pairs of electrodes located above and below the left eye and at the outer canthi of both eyes for the recording of vertical and horizontal eye movements, respectively. Bipolar electromyogram from the right masseter was also recorded for the purpose of artifact detection.

After MEG data acquisition, participants underwent MRI scanning with a 1.5T Philips Intera system for further reconstruction of the cortical surface.

### 2.5 MEG preprocessing

Raw MEG data were first processed to remove biological artifacts and other environmental magnetic sources that originated outside the head using the temporal signal-space separation method (tSSS) (Taulu et al., 2005) embedded in the MaxFilter program (Elekta Neuromag software). Data were converted to a standard head position (x = 0 mm; y = 0 mm; z = 45 mm). Static bad channels were detected and excluded from further processing steps.

Artifact correction caused by the vertical and horizontal eye movements, eyeblinks and R-R heart artifacts was performed on continuous data in Brainstorm (http://neuroimage.usc.edu/brainstorm; (Tadel et al., 2011) using the SSP algorithm (Tesche et al., 1995; Uusitalo and Ilmoniemi, 1997).

Data from two passive blocks were divided into 1610 ms epochs (from −610 ms to 1000 ms relative to the UP). Epochs with increased muscle activity contribution were excluded by thresholding the mean absolute signal values filtered above 60 Hz from each channel below 5 standard deviations of the across-channel average. After rejection of artifact-contaminated epochs, the average number of epochs taken into analysis was 183 ± 21 and 182 ± 21 for APW and NPW stimuli, respectively, before learning, and 181 ± 21 and 182 ± 20 for the same stimuli after learning (M ± SD). For all analyses reported below, baseline correction was computed using the interval from −210 to 0 ms before the stimulus onset (i.e., −610 – −410 ms relative to the UP).

### 2.6 Data analysis

Analyses were performed in two major steps. First, in the search for a general familiarization effect for the novel word-forms, the cortical responses to APW and NPW were compared between “before learning” and “after learning” conditions.

Secondly, we aimed to identify a putative semantic effect of pseudoword associative learning on neural activity elicited by pseudowords that acquired referential meaning. To this end, we analyzed the *APW* – *NPW* difference in time-locked responses before and after learning. We expected that while cortical responses to APW and NPW would not differ before learning, the differential response would emerge after learning because of acquired fine tuning of cortical representations toward the APW word-forms.

At each step, we analyzed MEG data both at the sensor and source levels in order to pinpoint the anticipated effects both in terms of their timing and involved cortical regions.

All further analyses were performed using MNE Python open-source software (http://www.nmr.mgh.harvard.edu/martinos) and custom-made scripts in Python.

#### 2.6.1 Sensor-level analysis

For sensor-level analysis, we took MEG signal from planar gradiometers that are known to attenuate signals from distant cortical sources and in effect behave as spatial high-pass filters (Vrba and Robinson, 2001; Garcés et al., 2017).

Large groups of sensors depicted in Figure 2 (*insert*) were chosen as ROIs for event-related filed (ERF) analysis, separately for the left and the right hemispheres. Each of the two ROIs included 31 pairs of gradiometers that covered frontal, temporal, and parietal selections of MEG sensors. Such wide ROIs at the sensor level were used on the basis of a large body of literature that demonstrated speech processing effects are mostly observed in wide perisylvian areas, including temporal, insular, inferior frontal, and inferior parietal cortices (Berwick et al., 2013; Hagoort, 2016; Hickok and Poeppel, 2016).

**Figure 2.**
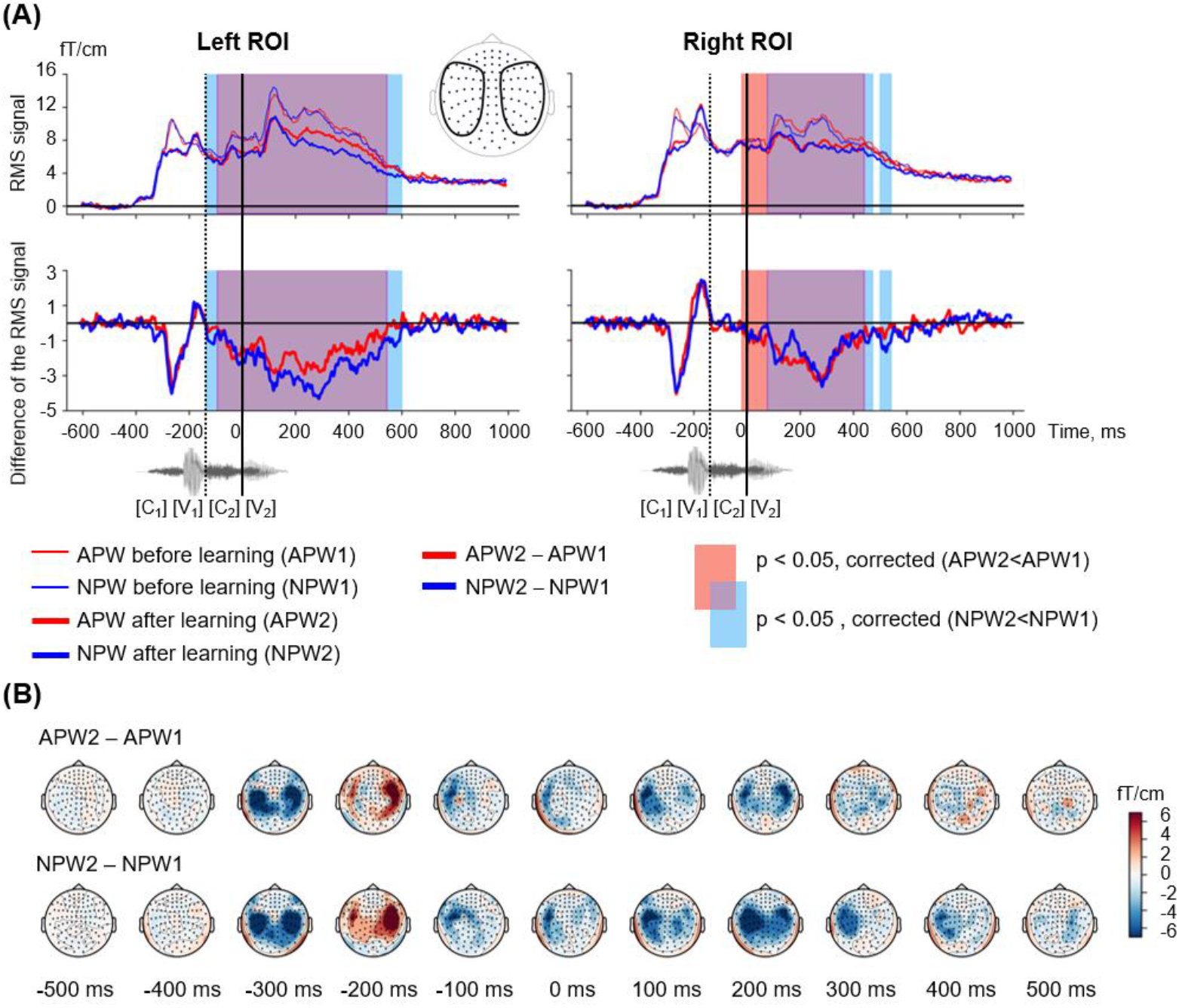
Familiarization effect in the sensor-space. **(A)** Time courses of the grand-average RMS signals (baseline-corrected). *Upper panel:* RMS time courses (baseline corrected) averaged over left and right ROIs (see insert) under passive listening to APW (red) and NPW (blue) stimuli presented before learning (thin red and blue lines) and after learning (thicker red and blue lines). *Bottom panel*: difference in RMS time courses between two passive listening blocks (*APW2* – *APW1* and *NPW2* – *NPW1*, thick red and blue lines, respectively). A significant repetition suppression effect for APW and NPW is shown as the blue and red shaded areas in RMS plots of APW and NPW trials, respectively (TFCE-based permutation statistics for “after learning” versus “before learning” contrasts); the purple shaded area corresponds to the temporal overlap of two effects. The waveform of the example stimulus ‘hicha’ aligned with the RMS timeline is shown at the bottom. Zero value on a timeline and a vertical solid line denote the onset of the fourth phoneme (word-form UP); a vertical dashed line shows the onset of the third phoneme. **(B)** Grand average topographic maps of the repetition effect magnitude for APW and NPW stimuli (*APW2* – *APW1* and *NPW2* – *NPW1* in the upper and bottom rows, respectively). Topographic maps are plotted in 100 ms steps; time is shown relative to the UP.

The data were first combined within each gradiometer pair by calculating the root-mean-square values (RMS) and then averaged across channel pairs; such averaging was performed independently within each of the two ROIs under each of the four experimental conditions (*APW1* and *NPW1* before learning, and *APW2* and *NPW2* after learning). The RMS signal was baseline-corrected using the interval from the −210 ms to 0 ms before the stimulus onset (−610 to −410 ms relative to the UP). A low-pass 6^th^-order Butterworth filter with a cutoff frequency 100 Hz was applied in order to smooth the RMS signals before statistical analyses; this procedure was done in order to improve the signal-to-noise ratio.

The RMS signals were compared for contrasts as specified below by using paired two-tailed t-test applied at each time point of the data within −410 to 1000 ms relative to the UP (0 – 1410 ms after stimulus onset). In order to provide correction for multiple comparisons, we applied the threshold-free cluster enhancement (TFCE)-based permutation statistical procedure; this approach takes into account both statistical intensity of each data point and its neighborhood via computing a “supporting area” for each data point (Mensen & Khatami, 2013). The permutation procedure involved 1,000 repetitions on surrogate data, which were generated from real data by swapping the two conditions for the entire time window in random subsets of participants. The TFCE-based permutation statistical procedure produced time intervals during which the timecourses significantly differed. The significance level was set at p < 0.05 (corrected).

#### 2.6.2 Source-level analysis

Individual structural MRIs were used to construct single-layer boundary-element models of cortical gray matter with a watershed segmentation algorithm (FreeSurfer 4.3 software; Martinos Center for Biomedical Imaging, Charlestown, MA, USA).

The cortical sources of the magnetic-evoked responses were reconstructed using distributed source modeling. Source estimation was performed using unsigned cortical surface-constrained L2-norm-based minimum norm estimation implemented in the MNE software suite. A grid spacing of 5 mm was used for dipole placement, which yielded 10,242 vertices per hemisphere. The ‘orientation constraint parameter’, which determines the extent, to which dipoles may deviate from the orthogonal orientation in relation to the cortical surface, was set to 0.4. Depth weighting with the order of 0.8 and the limit of 10 was applied.

For source space analyses, the MEG recording was downsampled to 200 samples per second; each new sample was calculated as an average of five adjacent timepoints for each vertex independently. Time window before stimulus onset (from −610 to −410 ms) was used as a baseline.

For distribution of reconstructed cortical sources over the two hemispheres, see Supplementary Figure.

Vertex-wise paired t-tests were performed for contrasts as specified below. If an effect was significant after correction for multiple comparisons at sensor level, we determined localization of the effect in the brain using the uncorrected significance threshold of p < 0.05 (see Gross et al., 2013).

### 2.7 Familiarization effects

#### 2.7.1 Sensor-level analysis

In order to reveal the time interval during which the familiarization effect was significant for both types of pseudowords, we contrasted RMS signals for “after learning” versus “before learning” conditions for APW and NPW pooled together, for each hemispheric ROI separately. Paired t-test with the TFCE permutation statistical procedure (see above for details) was applied at each time point of the entire RMS waveform (from −410 ms to 1000 ms relative to UP) to determine time intervals during which there was a significant difference between conditions.

For illustration purposes, we also repeated the same analysis separately for action pseudowords (*APW2* versus *APW1*) and non-action pseudowords (*NPW2* versus *NPW1*). Additionally, the differential topographic maps for ERFs elicited by APW and NPW stimuli separately were plotted in 100 ms steps (each plot representing data averaged across 35 ms).

#### 2.7.2 Source-level analysis

Cortical sources that exhibited the familiarization effect were reconstructed for time windows during which the effect was significant at the sensor level in the left and right ROIs (see above). The source-space data for APW and NPW types were averaged over these time intervals. We compared “before learning” and “after learning” conditions using vertex-wise paired t-test performed for two hemispheres (20,484 vertices). For the sake of comparison with previous passive word learning studies, we additionally repeated the same analysis using two successive intervals chosen roughly within the time window obtained at source level. The earlier interval (50–150 ms after UP) was taken because it was previously reported to demonstrate the ultra-rapid effects of word learning and discrimination (Shtyrov et al., 2010; Shtyrov, 2011; MacGregor et al., 2012; Kimppa et al., 2015). The later interval (150–400 ms) was taken as the timing of the significant semantic learning effect (see below). For illustrative purposes, the same analysis was repeated for APW and NPW trials separately.

### 2.8 Semantic learning effects

#### 2.8.1 Sensor-level analysis

We sought to identify the semantic learning effect by analyzing the contrast between cortical responses to APW and NPW types before and after learning. First, we checked for the significance of the *APW* – *NPW* difference for each of the two conditions (“before learning” and “after learning”) separately. Paired t-test with the TFCE permutation statistical procedure (see above for details) was applied at each time point of the entire RMS waveform (from −410 ms to 1000 ms relative to UP) to determine time intervals during which there was a significant difference between APW and NPW. We expected to find no statistically significant differences before learning and to reveal statistical effects after learning.

Next, we performed statistical analysis for the change in the difference between APW and NPW responses produced by the learning procedure, i.e. for the contrast [*APW2* – *NPW2*] versus [*APW1* – *NPW1*]. Where *APW1* and *APW2* stand for ERF time courses to passive presentation of APWs “before learning” and “after learning”, respectively, while *NPW1* and *NPW2* designate the responses to NPWs under the same two experimental conditions. Paired t-test with the TFCE permutation statistical procedure was used as described above.

To visualize the direction and dynamics of the effect, we plotted ERF topographic maps for the *APW* – *NPW* difference before and after learning at 100 ms steps (each plot representing data averaged across 35 ms).

#### 2.8.2 Source-level analysis

To reveal cortical regions, activation of which contributed to the “semantic learning” effect, the cortical sources of the effect were reconstructed within time intervals specified above (50–150 and 150-400 ms after UP). The former interval was taken with exploratory purposes for comparison with previous studies (see above), while the latter interval was taken on the basis of sensor level analysis (see above). We used a vertex-wise paired two-tailed t-test in order to contrast cortical activity averaged across the whole time interval for *APW1* versus *NPW1* (i.e. under “before learning” condition) and for *APW2* versus *NPW2* (i.e. under “after learning” condition).

Finally, we performed the most essential analysis: in order to explore the temporal dynamics of the semantic learning effect, we used a vertex-wise paired two-tailed t-test in order to contrast cortical activity for [*APW1* – *NPW1*] versus [*APW2* – *NPW2*] differences between conditions. This was done at the time points corresponding to the lowest p-values (see Gross et al., 2013) of the effect within the time interval that was identified at the sensor level. We averaged the source strength over 35-ms time intervals centered on the respective time points and considered only large cortical clusters including more than 20 adjacent vertices that demonstrated above-threshold significant effect at the respective time points. We then reconstructed activation timecourses for the obtained clusters of vertices.

## 3 Results

During the experiment, participants were presented with eight pseudowords (Table 1; Figure 1). The active task performed by participants was to learn specific associations between action pseudowords (APW) and motor actions by their hands and feet, while refraining from any responses to non-action pseudowords (NPW; Table 1). MEG was recorded during ‘Passive block 1’, which preceded wordform learning, and during ‘Passive block 2’, which followed learning (Figure 1B).

First, we examined possible general neural mechanisms related to familiarization with pseudoword word-forms. Second, we explored the cortical plasticity signature in the semantic brain network. We did not focus our analysis on pre-specified cortical regions or the time intervals of cortical responses, instead we employed an unbiased data-driven search (with correction for multiple comparisons) to reveal when and where in the cortex learning of novel word-forms and/or acquiring their semantics would induce neural activity changes.

### 3.1 Behavioral performance

All participants were successful with the task: average accuracy during the active performance block was 95.2 ± 5.8% (M ± SD), APW and NPW trials collapsed. Average d’ was 5.4 ± 1.1 (M ± SD). The total number of errors committed by participants during the active performance block was between 0 and 21 out of 320 trials.

### 3.2 Familiarization effects

#### 3.2.1 Sensor-level analysis

Figure 2A shows the root mean square (RMS) waveforms, calculated across gradiometers within left- and right-hemispheric regions of interest (ROIs), for passive presentation of APW and NPW in “before learning” and “after learning” conditions. These data illustrate the time courses of the overall signal strength of event-related fields (ERFs).

TFCE-based permutational statistical analysis revealed significant prolonged neural activity attenuation during the second passive block compared with the first one, and this modulation affected responses to both APW and NPW stimuli (Figure 2A). Response suppression significant for APW and NPW trials pooled together lasted from approximately −135 ms to 595 ms in the left ROI and from −20 ms to 550 ms in the right ROI – i.e., it started before the uniqueness point (UP) and lasted long after it. As can be seen in Figure 2A, the time intervals of significant suppression obtained for APW and NPW independently substantially overlapped.

Topographical maps (Figure 2B) demonstrated that this prolonged effect started approximately 100 ms before the UP, reached its maximum at 200 ms after the UP, and continued during the subsequent 300 ms gradually fading). Thus, the second block of passive presentations of the same word-forms following learning led to a bilateral neural response attenuation to the temporal combination of the successive phonemes for both APW and NPW stimuli. A relatively early onset along the RMS time course suggests that the response attenuation was likely linked to the onset of the third rather than the fourth phoneme during auditory word-form processing.

#### 3.2.2 Source-level analysis

To identify cortical areas that contributed to neural repetition suppression resulting from familiarization with the novel pseudowords, we analyzed the data in the source-space. We evaluated cortical clusters that underwent significant suppression across the whole-time interval revealed by RMS analysis for APW and NPW trials pooled together (Figure 3A). Additionally, we examined two consecutive time periods within the interval that was significant at the sensor level: earlier (50–150 ms after the UP) and later (150–400 ms after the UP) ones (Figure 3B-D). The source-space analysis showed that significant neural activity suppression occurred within both time periods and affected widely distributed cortical areas in both hemispheres, including the lateral and opercular surface of the temporal lobe, insula, lateral and ventral parts of the motor cortex, and inferior parietal regions (Figure 3).

**Figure 3.**
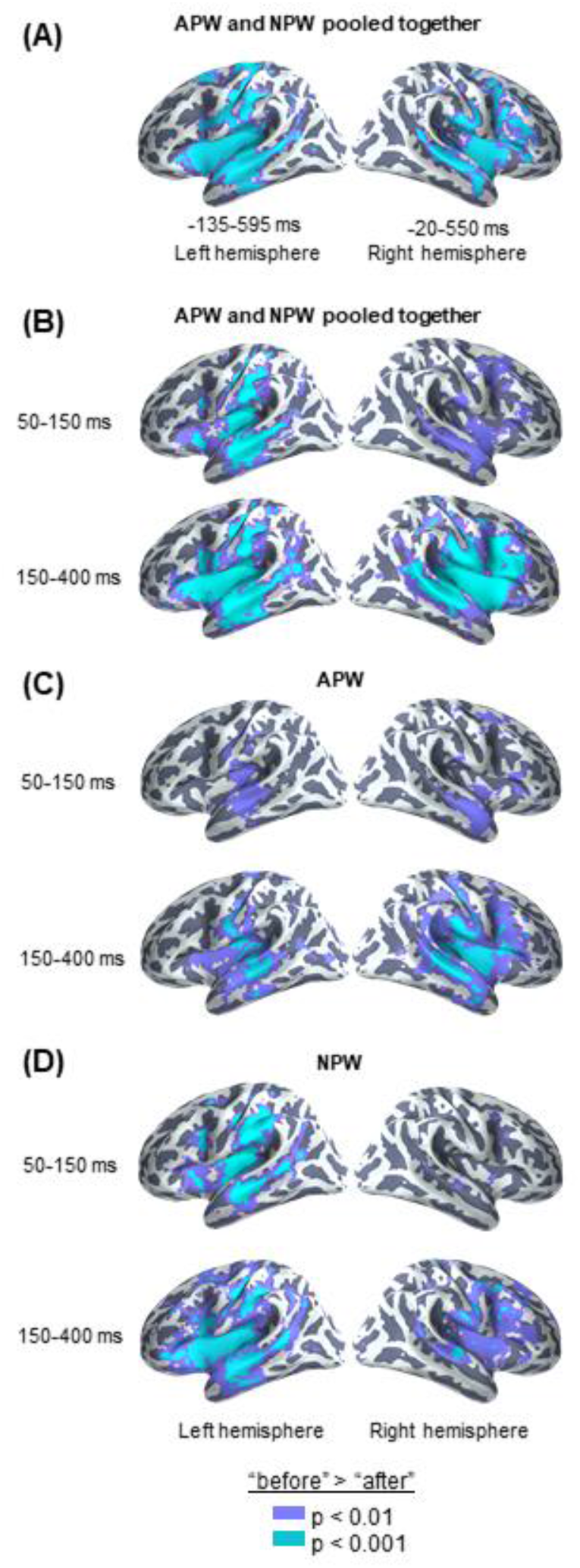
Familiarization effect in the source-space. Statistically thresholded maps (voxel-wise paired t-test, p < 0.01 and p < 0.001, are shown in purple and light-blue colors, respectively) for “after learning” versus “before learning” contrasts. **(A)** Analysis performed on data averaged across the time intervals revealed by the RMS analysis at sensor level (- 135 – 595 and −20 – 550 ms in relation to the UP, for the left and the right hemispheres respectively), with the APW and the NPW conditions pooled together. **(B-D)** Analyses performed on data averaged across two time windows: 50–150 ms after the UP and 150–400 ms after the UP: **(B)** APW and the NPW trials collapsed, **(C)** APW analyzed separately, and **(D)** NPW analyzed separately, respectively.

### 3.3 Semantic learning effects

#### 3.3.1 Sensor-level analysis

Both RMS signal timecourses (Figure 4A) and ERF topographical maps (Figure 4B upper row) demonstrated that whereas cortical activity evoked by the two pseudoword types were almost indistinguishable before learning, the strength of the differential neural responses to APW significantly increased after the learning procedure in the left ROI (Figure 4B, middle row). Indeed, analysis of the significance of the *APW- NPW* difference separately before learning and after learning using TFCE-based permutational procedure revealed that neural responses to APW and NPW stimuli did not differ statistically before learning (Figure 4A), while after learning they were statistically different within a prolonged time window from 145 to 615 ms after the UP in the left hemisphere (Figure 4A, left panel). No statistical significance was found in the right ROI (Figure 4A, right panel). Remarkably, unlike familiarization-related changes in the neural response, the semantic-sensitive transformation of cortical activity was apparently bound to the moment in time when the two pseudoword types started to be recognizable by the brain.

**Figure 4.**
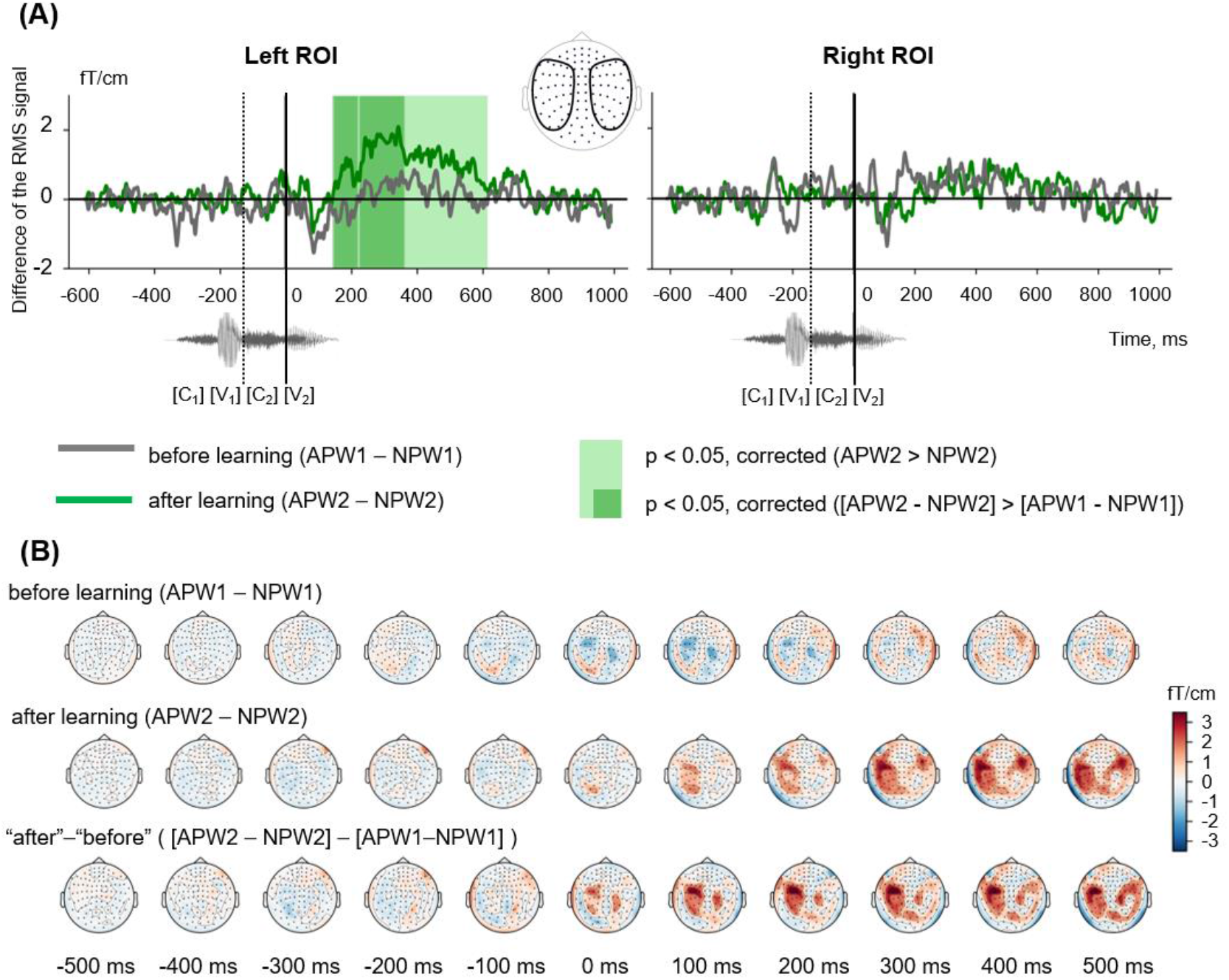
Semantic learning effect in the sensor-space. **(A)** Grand average differential RMS time courses (baseline-corrected) for APW versus NPW contrast “before learning” (*APW1* – *NPW1*, grey) and “after learning” (*APW2* – *NPW2*, green). The light green shaded area marks the time interval in “after learning” neural responses that correspond to a significant *APW2* > *NPW2* contrast according to TFCE permutation statistics; no such significant difference was found for “before learning” condition. The dark green shaded areas designate the time intervals for a significant differential learning effect ([*APW2* – *NPW2*] > [*APW1 –* APW1]). A zero value on a timeline and a vertical solid line denote the onset of the fourth phoneme in the auditory pseudowords (wordform UP); a vertical dashed line shows the onset of the third phoneme. **(B)** Grand average topographic maps of differential ERF for APW versus NPW contrast “before learning” (*APW1* – *NPW1*, top row) and “after learning” (*APW2* – *NPW2*, middle row). The bottom row represents the differential learning effect: “after leaning” minus “before learning” ([*APW2* – *APW2*] > [*APW1* – *APW1*]). Topographic maps are plotted in 100 ms steps; time is shown relative to the UP.

Next, we checked for the significance of the change in the difference between APW and NPW responses produced by the learning procedure ([*APW2* – *NPW2*] versus [*APW1* – *NPW1*]). The same TFCE-based analysis of these differential RMS signals produced two statistically significant intervals for the associative learning effect in the left ROI: 144–217 ms and 226–362 ms after the UP (Figure 4A, left panel, and Figure 4B, bottom row). Since the gap that separates the two temporal clusters was very narrow (9 ms), we merged them. Statistical significance maxima of the differential effect along the RMS timecourse were observed at 190, 265, and 325 ms after the UP. Again, no statistically significant effect was found in the right ROI.

#### 3.3.2 Source-level analysis

Proceeding from the sensor-level results presented above, we chose to evaluate the semantic learning effect only in the left ROI within the time interval of 150–400 ms after the UP; thus, the following source reconstruction was restricted to the left hemispheric cortical responses within this time interval. For each of the four conditions, the source strength was averaged across the above interval before the statistical comparisons. Figure 5 (right panel) demonstrates the after-learning enhancement in the activation strength of cortical sources in response to APW compared with NPW within “before learning” and “after learning” conditions (passive blocks 1 and 2 respectively). The largest contribution to the effect was from anterior parts of the superior temporal sulcus (aSTS)/middle temporal gyrus (MTG), insula/frontal operculum, triangular portion of inferior frontal gyrus (IFG), and the orbital area of prefrontal cortex. The 50–150 ms time interval was also analyzed for the sake of comparison with previous studies: during this earlier interval, virtually no identifiable activation changes caused by learning could be seen (Figure 5, left panel).

**Fig. 5.**
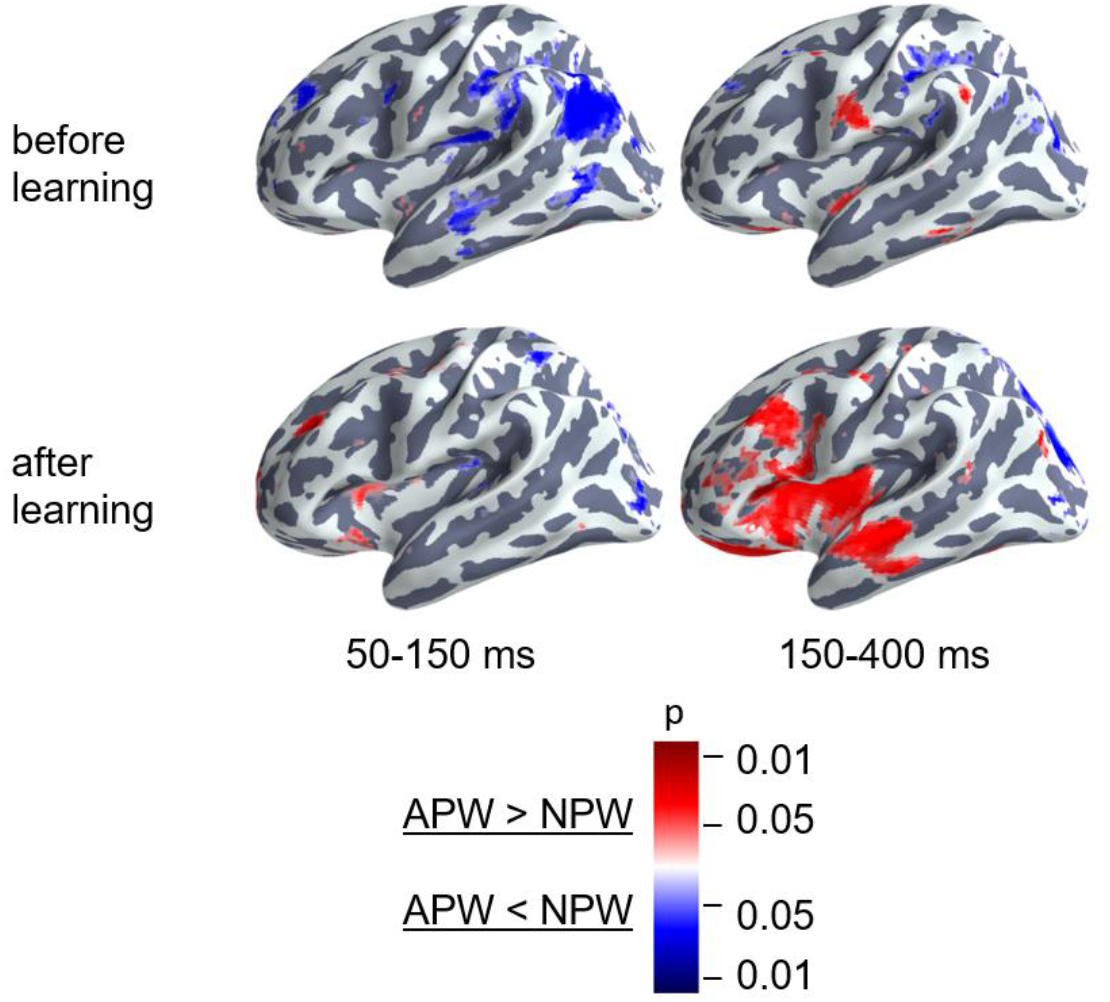
Cortical areas engaged in the semantic learning effect in the source space. Statistically thresholded cortical topography for the responses to APW versus NPW “before learning” (*APW1* – *NPW1*, top row) and “after learning” (*APW2* – *NPW2*, bottom row) (voxel-wise paired t-test, p < 0.05, uncorrected). The 150–400 ms time interval corresponds to the significant learning effect on APW versus NPW contrast according to the RMS data, which survived after correction for multiple comparisons (right column). The 50–150 ms time interval (left column) is presented for the sake of comparison with previous studies. The color scale represents p-values; color denotes the sign of the effect: red for APW > NPW and blue for APW < NPW.

To explore the temporal dynamics of the semantic learning effect further, we reconstructed the cortical sources within 35-ms time frames centered on time points that corresponded to statistical significance maxima of the differential effect along the RMS timecourse: 190, 265, and 325 ms after the UP. Voxel-wise statistical comparisons between “after leaning” (*APW2* – *NPW2*) versus “before learning” (*APW1* – *NPW1*) were performed for each of three time frames independently. Significant clusters were defined as groups of neighboring significant voxels (p < 0.05, uncorrected); see Table 2 for the list of the respective clusters.

**Table 2.** Brain regions involved in “semantic” learning.

| Cluster localization | The most significant vertex within each cluster |  |  |  |  |
| --- | --- | --- | --- | --- | --- |
|  | MNI coordinates<br>(x, y, z) |  |  | T-value | p-value<br>(uncorrected) |
| **Ventral premotor (VPM) and opercular part of inferior frontal gyrus (IFG)**<br>**190 ms** | -52.64 | 20.44 | 17.64 | -3.97 | 0.001 |
| **Insula and frontal operculum**<br>**265 ms** | -39.88 | 2.53 | 11.42 | -3.76 | 0.001 |
| **Triangular and orbital inferior frontal gyrus (IFG)**<br>**325 ms** | -46.18 | 25.96 | 11.59 | -3.33 | 0.003 |
| **Intraparietal sulcus (IPS)**<br>**190 ms** | -39.19 | -43.38 | 37.83 | -3.22 | 0.001 |
| **Anterior superior temporal sulcus (aSTS)**<br>**265 ms** | -46.45 | -17.63 | -12.15 | -3.30 | 0.003 |
| **Temporal pole (TP)**<br>**325 ms** | -45.04 | 4.83 | -25.58 | -3.74 | 0.001 |

Figure 6 shows the cortical location of the clusters of vertices reconstructed at each of the three sequential time frames, as well as their activation timecourses before and after learning. Initially, around 190 ms post-UP, a learning-related selective response to APW emerged in cortical areas surrounding the Sylvian fissure: aSTS, ventral premotor cortex, and the anterior part of intraparietal sulcus and insula. Once it appeared, differential activation in these areas was mostly sustained until response termination. After ~250 ms, activation spread to more anterior brain regions, and by 325 ms after the UP, it reached the pole of the left temporal lobe and the triangular part of the left IFG extending to its orbital part. Thus, the spatiotemporal pattern of semantic-learning-related neural activity in our study was generally consistent with the current hierarchical models of auditory word processing that imply the presence of an anterior-directed stream of word-recognition pathways (Hagoort, 2016; Hickok and Poeppel, 2016).

**Figure 6.**
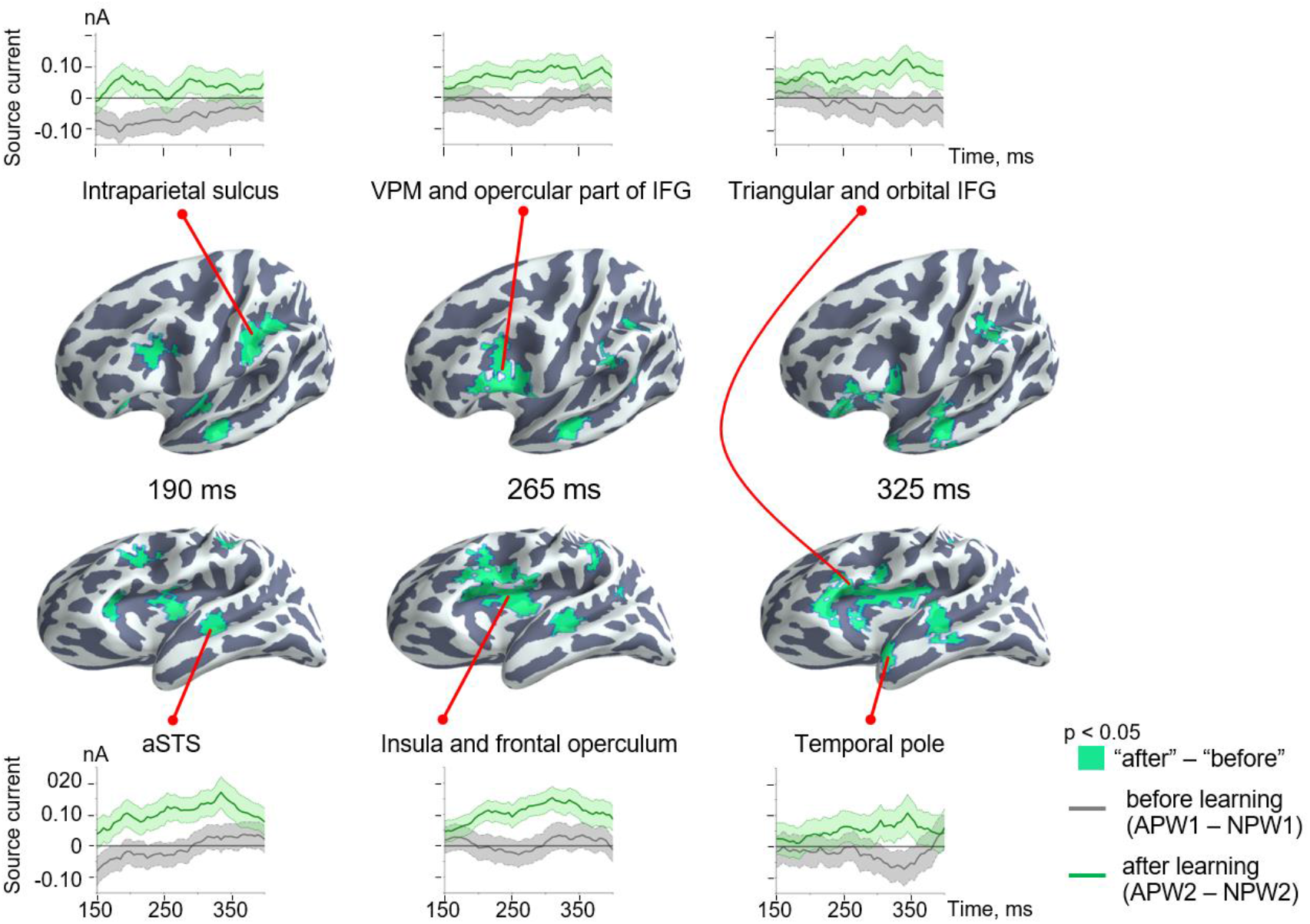
Spatio-temporal dynamics of the semantic learning effect in the source-space. Clusters are displayed on the cortical surface of the inflated left hemispheres shown at two different angles of view in order to represent deep locations within the Sylvian fissure. The timecourses at the top and at the bottom represent grand-averaged differential response strength for the clusters of cortical vertices across time for “before learning” (*APW1* – *NPW1*, gray lines) and “after learning” (*APW2* – *NPW2*, green lines) conditions. Shaded areas on timecourses represent standard errors.

## 4 Discussion

### 4.1 Overview

Whether short-term learning of new words can induce rapid changes in cortical areas involved in distributed neural representation of the lexicon is a hotly debated topic in the literature. To answer this question, we examined the MEG time-locked responses elicited in the cerebral cortex by passive presentation of eight novel pseudowords before and after an operant conditioning task. The task forced the participants to perform an active search for word-form meaning, as four unique wordforms acquired meaning that referred to movements of participants’ body parts in a way similar to real action words (action pseudowords – APW) and the other four word-forms were not associated with any particular meaning (non-action pseudowords – NPW). By comparing learning effects between the two types of pseudoword stimuli, either with learnt semantics or without it (while both types were equally well familiarized through an equal number of presentations during the experiment), we expected to observe the emerging cortical signature of newly learned meaningful words.

There were three main findings in the current study. Deep familiarization with both APW and NPW acoustic word-forms led to a highly reliable and long-lasting suppression of cortical responses starting before or around the UP in both hemispheres. Semantization of the new word-forms was followed by a relative learning-related increase in cortical activity to meaningful word-forms (APW) compared with meaningless ones (NPW) at around 150–400 ms after UP, which was lateralized to the left hemisphere. These learning-related changes in left-hemispheric cortical responses to semantically meaningful words were localized to the perisylvian cortex starting at ~ 150 ms, and to the higher-tier speech areas (temporal pole and triangular/orbital part of inferior frontal sulcus/gyrus) starting after ~250 ms from the word-form UP. All of these rapid learning effects were observed during passive presentations of the pseudowords that followed successful learning (greater than 90% accuracy) and repetitive performance of actions implied by the meaning of the newly learned words.

### 4.2 Familiarization effects

Our finding of a strong and highly reliable repetition suppression effect in the time-locked response to repeated passive presentation of pseudowords (both APW and NPW) stands in stark contrast with the previously reported EEG/MEG findings, according to which repetition suppression was characteristic for real words, while for pseudowords repetition caused the inverse effect, namely response enhancement (Shtyrov et al., 2010; Shtyrov, 2011; Kimppa et al., 2015). Notably, the repetition suppression effect in our data remained significant even we investigated exactly the same time interval, 50–150 ms after the word-form UP, which was previously reported to contain an enhanced evoked response to repeated pseudowords (Figure 4). How can stimulus repetitions have opposite effects on cortical responses depending on the way the stimuli were manipulated within the recent experience between the successive presentations?

Most probably, a degree of word-form familiarization, which might be collectively greater in the current experiment compared with the previous ones, is critically important for the sign of the neural repetition effect. The effect of repetition suppression is ubiquitous in the brain and well described for various sensory modalities (Grill-Spector et al., 2006; Gotts et al., 2012). Neural response reductions within a one-session stimulus repetition is thought to be indicative of familiarization memory traces, which scales down the neural representation of the stimulus without sharpening it (McMahon and Olson, 2007; Weiner et al., 2010), for review see Gotts et al. (2012). While “repetition-enhanced neural responses” were reported less frequently in the human literature, they are predominantly characteristic of the repetition dynamics for unfamiliar stimuli (Henson et al., 2000) or for those with poor perceptibility (Turk-Browne et al., 2007). Moreover, as demonstrated for a visual modality, repetition effects for unfamiliar stimuli can turn from enhancement to suppression when the number of stimulus repetitions increases, a phenomenon that possibly reflects a shift in neuronal responses depending on the degree of stimulus familiarity and on-line accessibility of its neuronal representation (Müller et al., 2013). Since the cumulative number of repetitions for each pseudoword in our experiments (between 150 and 200, depending upon individual learning rate) did not differ much from that used during passive presentations in the previous studies (about 150-200), the opposite sign of repetition effects could hardly result from just a different number of stimulus repetitions. Yet, to continue this logic, another possibility is that deep familiarization with APW and NPW word-forms during our operant conditioning procedure completely changed the repetition effect: instead of increasing neural responses to previously unfamiliar word-forms, it decreased them when the word-forms became well-recognized concatenations of phonemes. Indeed, although we observed transient repetition-related changes in time-locked ERF components elicited by auditory word onset well before the UP (Figure 2), a long-lasting and highly reliable attenuation of time-locked activity occurred approximately 300 ms after stimulus onset, when the word-form began to be discriminable from each other. In fact, the onset of this reduction started even 100 ms earlier than UP, probably as a response to the appearance of the third phoneme in the word-form, which, unlike the UP, was not sufficient to distinguish all eight pseudowords, but rather allowed identification of the difference between APW–NPW pairs (see Methods for details).

The above considerations suggest that our findings of the strong suppression of neural responses to novel acoustic word-forms, which started to be familiar through the experimental procedure, most probably reflect a mechanism of familiarization memory. This mechanism is one of the components of the recognition memory system that is responsible for judging the prior occurrence of a stimulus based on detecting stimulus familiarity. It is thought to be centered on extra-hippocampal regions of the medial temporal memory system (Brown and Aggleton, 2001; Bird, 2017), and it is associated with repetition suppression of neural responses to a familiar stimulus in perirhinal and neocortical structures that appear in one-session experiment and then last over days (Brown and Aggleton, 2001). Synaptic depression plasticity in the perirhinal cortex seems to play a critical role both in the activity-dependent suppression of neural responses and visual recognition memory (Griffiths et al., 2008). Repetition-sensitive neuronal phenomena (either suppressive or enhancive) accompany perceptual learning, and although they are unlikely to be its main underlying neural cause, they still might represent one of its mechanisms (Gotts et al., 2012); also for different opinions see McMahon and Olson (2007).

While primarily determined by novelty/familiarity of a complex auditory stimulus, which is processed by the perirhinal cortex of the medial temporal lobe, repetition-sensitive neocortical responses are hardly indicative of learning-related neocortical plasticity. In other words, neither repetition suppression of time-locked responses to novel word-forms found in our experimental settings nor the repetition enhancement effect resulting from their passive presentation (Shtyrov et al., 2010; Shtyrov, 2011; Kimppa et al., 2015) can be considered true indicators of rapid cortical plasticity during word learning.

### 4.3 Semantic learning effects

In order to conclude that experience-dependent modification of neocortical activity during word learning complies with the criteria of rapid cortical plasticity, one should at least provide evidence that (1) cortical electrophysiological responses to unfamiliar word-forms are predictably and persistently modified by the experience obtained within a single experimental session, and index cortical plastic changes that lead to “experientially-induced tuning” toward a specific word-form neural representation, and (2) newly formed cortical representation is not only tuned to a particular concatenation of the phonemes but also possesses referential meaning, i.e., its activation is linked to increased activation of the “semantic network” that encompasses multisensory higher-tier speech areas involved in semantic associations (Davis and Gaskell, 2009). The contrast between neural responses elicited by action-associated (APW) and non-associated (APW) pseudowords before and after operant conditioning answered the question to what extent the learning-related neural dynamic complies with these criteria.

Animal neurophysiological findings evidence that while neural activity in the auditory cortex decreases overall with stimulus repetition, firing rates become more selectively tuned toward stimuli that attain behavioral relevance, and the neural cells that encode such stimuli may maintain their firing rate levels or decrease them much less than other cells (Weinberger, 2004; Blake et al., 2006; Kato et al., 2015). If improved stimulus selectivity and sharpening of neural representations did occur for APW, we would expect that after operant conditioning, cortical brain responses to APW would relatively increase compared with NPW. This finding is exactly what we observed while contrasting APW–NPW differences before and after learning (Figure 6).

Indeed, the only factor that affected the auditory perception of APW and NPW stimuli during the second passive presentation was their unique relatedness to a specific motor action in the prior active experimental blocks. Acoustical features across APW–NPW pairs were well counter-balanced within subjects across the eight pseudowords (see Methods), and neural responses to pseudowords of both types did not differ before learning (Figures 5 and 6). Additionally, our findings cannot be explained by differences in selective attention to or in familiarization with APW–NPW pairs during learning. The learning procedure itself did not introduce any bias toward APW word-forms, as it required the subject to attentively discriminate between both stimulus types, which were repeated the same number of times and interleaved into pseudorandom sequences. Even with respect to action-relatedness, both types of pseudowords required a similar level of perceptual decision-making activity, because a subject had to either commit a motor response to the APW stimuli or refrain from it for NPW ones. Despite having behavioral relevance, NPW word-forms lacked unique referential meaning to a specific event, a core property of lexical items in human language. Thus, our results on experiential modification of human cortical responses to neutral auditory stimuli through an operant conditioning association procedure bears a striking resemblance to that described in single-cell recordings in monkey auditory cortex (Blake et al., 2006; Kato et al., 2015). These data may be considered as some of the first convincing evidence of rapid cortical plasticity in human neocortex in relation to novel word learning.

Notably, while repetition suppression emerged rather early in the timecourse of the magnetic response to word-forms (possibly before the UP), the semantic learning effect for APW–NPW contrast occurred relatively late in the time-locked neural activity, starting about 150 ms after the word-form UP (Figures 3 and 5). At this point, the differential left-hemispheric response to APW encompassed mainly the aSTS, ventral premotor cortex, insula/opercular part of IFG, and anterior IPS. These left perisylvian regions are heavily interconnected through the classic arcuate fasciculus pathway that connects superior temporal regions with extended Broca’s area, but also through a parallel pathway that projects from the superior temporal sulcus (STS) to the inferior parietal region. These routes are thought to participate in acoustic/phonological transcoding (Catani et al., 2005). The recurrent motor-perceptual interaction is known to facilitate speech perception of unfamiliar speech stimuli, e.g., distorted speech and novel or low-frequency words (Wu et al., 2014; Stokes et al., 2019). Therefore, greater involvement of the entire perisylvian network into the APW compared to the NPW response in our experiment may indicate that newly learned semantic association boosted perceptual processing of incoming novel linguistic stimuli.

Our interpretation is generally in line with MMN results of Hawkins and colleagues, who described an increased MMN wave peak at 140 ms after the word UP in response to auditory pseudowords that acquired association with visual images (Hawkins et al., 2015). However, in our case, enhanced neural response to APW spanned from approximately 144 ms after the first meaningful phoneme and onwards, which clearly occurred later than the MMN wave. We speculate that rather than reflecting a rapidly detected phonological difference in the fourth phonemes between APW and NPW, the differential response to APW points to enhanced activity of neuronal circuitry that mediates sensitivity for the temporal sequence of the phonemes that corresponded to the APW word-forms. There is ample evidence in the literature on the existence of higher-level auditory neurons that contain the combinatorial code for the whole auditory word-form and operate approximately 150–250 ms after the moment when a word-form becomes identifiable (Brink et al., 2001; DeWitt and Rauschecker, 2012). In addition to later timing compared with the MMN, the APW differential response localization to the anterior superior temporal gyrus/superior temporal sulcus is compatible with its putative origin from higher-order combinatorial phonological representations of the entire word-form. Imaging studies localized processing of multisegmental word-forms to the left anterior STG/STS, downstream of the middle-posterior STS/STG, which underlies specific phoneme discrimination (Chang et al., 2010; DeWitt and Rauschecker, 2012). The strict left lateralization of the APW response (Figure 4) is also concordant with the putative site of auditory word-form recognition (see meta-analysis by DeWitt and Rauschecker, 2012).

Therefore, our findings suggest that the tuning of higher-order combination-sensitive neurons in aSTS for a word-form is contingent upon experience of its unique action relevance obtained within one experimental session. In other words, even short-term active search for auditory-action association, or effortful semantization of word-form provided by our experimental settings, facilitates or even triggers strengthening of the cortical network that underlies the phonological aspect of lexicality: lexical representation of the respective coherent word-form.

The aSTS-centered cortical network, which is thought to contain lexical representations of real-word word-forms, does not store semantic information itself, but rather it interfaces with the semantic network that is widely distributed across the brain (DeWitt and Rauschecker, 2012). The question as to whether prolonged post-stimulus enhancement of neural responses to pseudowords (Figure 4) reflects facilitated activation of features of the long-term memory representations that were briskly associated with a new lexical item. Our data may provide a tentative answer to this question. APW-related differential activation timecourses (Figure 6) suggest that after 200–250 ms, differential activation spreads from the perisylvian cortex toward more anterior cortical areas along both ventral and dorsal speech processing pathways (Romanski et al., 1999; Rauschecker and Tian, 2000). Specifically, the activation timecourses in the ventral speech stream point to the later involvement of areas identified anatomically as the temporal pole, which was previously implicated in the semantic access (Binder and Desai, 2011; Ralph et al., 2016). Concurrently, relatively delayed activation in the dorsal stream occurs in the triangular part of the IFG that encompasses the classical Broca’s area as well as IFG orbital part, i.e., the left ventrolateral prefrontal cortex, thought to subserve controlled semantic retrieval (Thompson-Schill et al., 1997; Ralph et al., 2016). Localization of the late portion of the APW response to higher-order “semantic” cortical areas assumes that after learning, APWs selectively increase activity of the semantic network, i.e., attain the eminent property of the time-locked cortical responses in the N400 range to real words compared with pseudowords (Cheng et al., 2014).

Notably, the activity of the left-hemispheric aSTS and ventral premotor speech areas involved in phonological processing of the auditory word-form persisted throughout the entire APW differential response approximately from 190 until 350 ms after the UP (Figure 6). This long-lasting activation of the phonological word-form representations is consistent with the principle of refining processing of complex stimulus features in hierarchical reentrant system (Bullier, 2001; Di Lollo, 2012) and may reflect recurrent interaction between different hierarchical levels of auditory word-form analysis. It is generally assumed that the main function of re-entrant signals is modulatory, and they may prolong and modify activity induced by bottom-up signals by way of integrating neuronal responses at each level of the pathway under the top-down influence from the higher order areas. From this view, we assume that while the early differential activity in the perisylvian areas appears to be stimulus-driven, the later activity there presumably depends on top-down signaling from higher-order speech cortical areas involved in semantic retrieval.

### 4.4 Conclusions

In summary, we would argue that according to criteria proposed by Davis and Gaskell (2009), our data evidence that cortical representations of both phonology and semantics of previously unfamiliar words may be formed following 1–2 hours of active associative learning. Particularly, we observed word learning effects in the higher-tier cortical areas that underlie semantic processing of real words. Such changes were in fact observed in these regions after an overnight consolidation (Davis and Gaskell, 2009), yet we observed these effects within a single experimental day.

This conclusion raises the question as to why a rapid cortical activity modulation by a newly learned word would be found in our MEG study, while the blood-oxygen-level-dependent (BOLD) response of cortical areas consistently remain largely unaffected during the hours after associative learning (Davis and Gaskell, 2009). There are two putative explanations for this discrepancy. First, the discordance between MEG and fMRI findings may result from different modes of neural activation captured by changes in an evoked, time-locked response in the MEG and BOLD signals. Given that the BOLD signal integrates brain hemodynamic changes over several seconds, short-lived and synchronized neural activation that contributes to MEG/EEG time-locked response could be difficult to detect with fMRI (Engell et al., 2012). Thus, rapid formation of cortical representations of a newly acquired word may increase time-locked yet transient cortical activation elicited by passive presentations of these stimuli; such coherent cortical activation is reflected in enhanced MEG time-locked ERFs. Only further strengthening of plastic cortical changes during a consolidation process might make them detectable using the fMRI recording technique.

Another explanation, which is not necessarily mutually exclusive with the first one, focuses on the difference in the associative learning procedure between our study and the previous fMRI research. The latter studies tested the involvement of cortical structures in adult experience-dependent neuroplasticity using paired-associative learning between auditory pseudowords and visual images. However, the findings of Blake and colleagues (Blake et al., 2006) demonstrated that the learning-induced increase in response selectivity of auditory neurons is observed following an active operant conditioning, but not after passive reward-based associative learning. As the authors suggested, the successful reward-association plasticity that results from operant conditioning might be related to a greater involvement of neuromodulatory brain systems triggered by an increase in the subject’s motivation for active search for stimulus-action pairing. Indeed, subjects’ activity in terms of “active selection”, “active discovery” and “repeated retrieval” has recently been ascertained as a prerequisite condition of rapid cortical plasticity – for word semantic learning and other forms of learning (Hebscher et al., 2019). In other words, whether a new word will be learned depends on personal engagement into the learning process, wisdom that ages ago was recognized by psychological science: “Student engagement is the product of motivation and active learning. It is a product rather than a sum because it will not occur if either element is missing” (Barkley, 2009).

## 7 Acknowledgments

We wish to thank Alena A. Zhukova for involvement in the initial data analysis, Platon K. Pronko for valuable comments on data analysis, and Alexey N. Kozlov for manufacturing the pedals.

