## Supplementary Figure for "Rapid cortical plasticity induced by active associative learning of novel words in human adults"

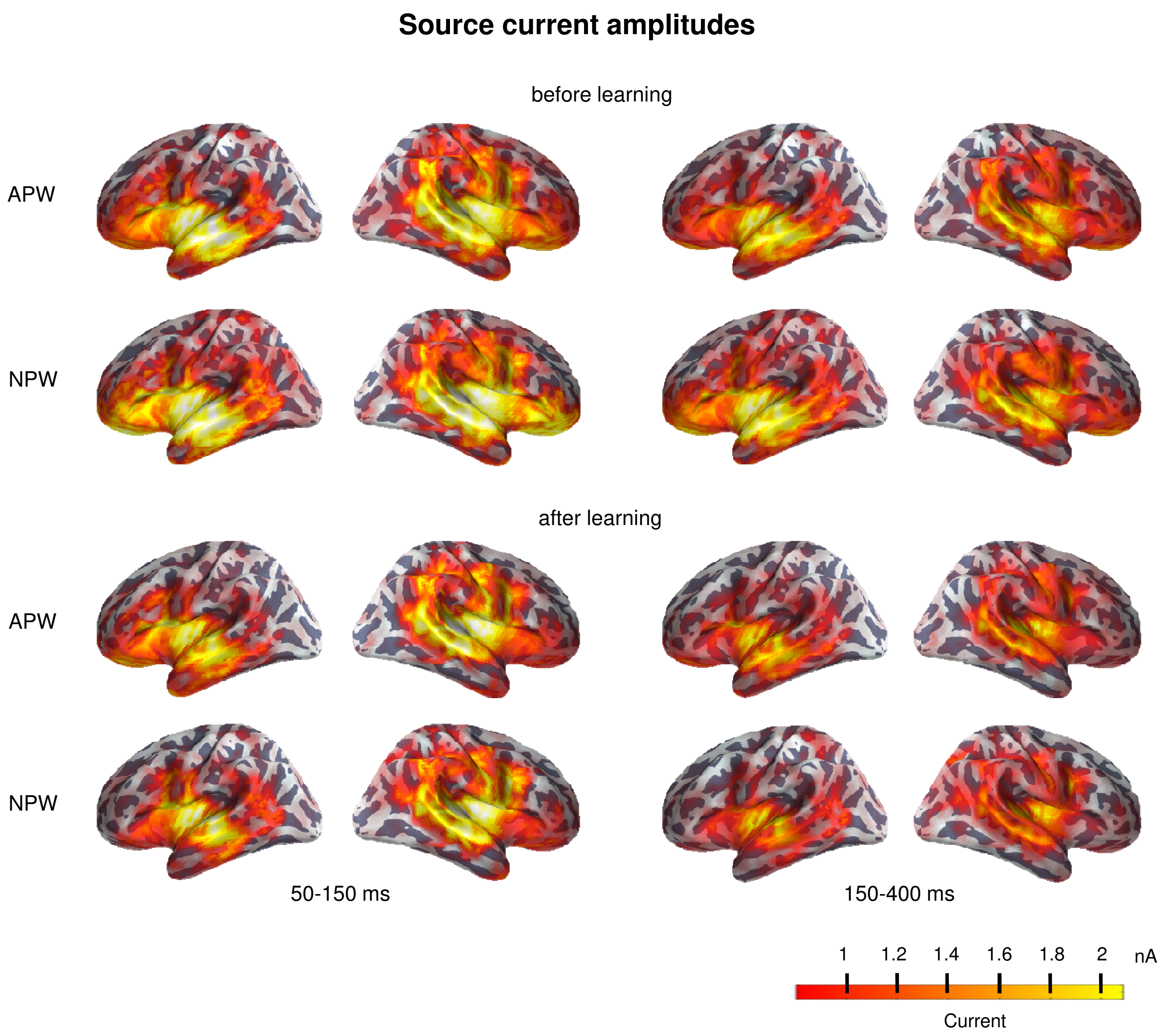


**Supplementary figure**. Reconstructed cortical sources over the hemispheres for APW and NPW before and after learning.
